# A dark matter in *sake* brewing: Origin of microbes producing a *Kimoto*-style fermentation starter

**DOI:** 10.1101/2022.11.28.518188

**Authors:** Kohei Ito, Ryo Niwa, Ken Kobayashi, Tomoyuki Nakagawa, Genki Hoshino, Yuji Tsuchida

**Affiliations:** BIOTA Inc., Tokyo, 101-0022, Japan; Graduate School of Medicine, Kyoto University, Kyoto, 606-8501, Japan; Faculty of Applied Biological Sciences, Gifu University, Gifu, 501-1193, Japan; Tsuchida Sake Brewing Company, Gunma, 378-0102, Japan

**Author notes:** Equal contribution (Kohei Ito, Ryo Niwa). **Correspondence:** Kohei Ito.

**Keywords:** Japanese *Sake*, Microbial fermentation, *Kuratsuki*, Microbiome, *Kimoto*-style

## Abstract

In *Kimoto*-style fermentation, a fermentation starter is produced before the primary brewing process to stabilize fermentation. Nitrate-reducing bacteria, mainly derived from brewing water, produce nitrite, and lactic acid bacteria such as *Leuconostoc* can proliferate because of their tolerance towards low temperature and their low nutritional requirements. Later, *Lactobacillus* becomes the dominant genus, leading to weakly acidic conditions that contribute to control yeasts and undesired bacterial contaminants. However, the sources of these microorganisms that play a pivotal role in *Sake* brewing have not yet been revealed. Thus, comprehensive elucidation of the microbiome is necessary. In this study, we performed 16S rRNA amplicon sequencing analysis after sampling from floor, equipment surfaces, and raw materials for making fermentation starters, including *koji*, and water in *Tsuchida Sake* brewery, Gunma, Japan. Amplicon sequence variants (ASVs) between the external environments and the fermentation starter were compared, and it was verified that the microorganisms in the external environments, such as built environments, equipment surfaces, and raw materials in the sake brewery, were introduced into the fermentation starter. Furthermore, various adventitious microbes present in the fermentation starter of early days and from the external environments were detected in a nonnegligible proportion in the starter, which may impact the taste and flavor. These findings illuminate the uncharacterized microbial dark matter of sake brewing, the sources of microbes in *Kimoto*-style fermentation.

## 1. Introduction

### Description of microbial transition of a fermentation starter of *Kimoto*-style *Sake*

Fermentation is an ancient biotechnology that has been used to improve the safety, storage capacity, flavor, and nutrition of food. The most modern fermented foods are manufactured by adding fermentation starters for improving stability and reproducibility. In contrast, the fermentation process has traditionally been carried out by utilizing microorganisms from raw materials, starter cultures, and the environment around the fermentation batches. Among fermented foods, *Sake* is the traditional alcoholic beverage of Japan. *Sake* is made from rice through starch saccharification by *Aspergillus oryzae* and alcoholic fermentation by *Saccharomyces cerevisiae*. The beverage is traditionally brewed using a fermentation starter called *Moto*.

*Moto* is a yeast mash made from a nutritious mixture of rice, *Koji* and water. In the preliminary stage of the primary brewing process, *Moto* is manufactured to prevent the proliferation of undesired contaminants and to promote smooth alcoholic fermentation. Based on the manufacturing methods of *Moto, Sake* is divided into three types, *Sokujo, Kimoto*, and *Yamahai. Sokujo-style Moto* is a modern method to add food-grade quality lactic acid and decrease pH before alcoholic fermentation by yeast. Lactic acid inhibits contaminations of unintended yeasts and bacteria from external environments into the fermentation starter. Since *Sokujo* relies on lactic acid to control the pH of *Moto*, a desirable condition can be created relatively quickly, leading to stable production of the starter. Meanwhile, the *Kimoto*-style *Moto* is a traditional preparation method for the starter culture and is manufactured by inducing the growth of lactic acid bacteria properly. Lactic acid bacteria produce lactic acid, which contributes to creating an environment conducive to the growth of yeast. The *Yamahai* style is similar to the *Kimoto* style but made without grinding rice. *Koji*, mold (*Aspergillus oryzae*) on rice, degrades solid rice and initiates alcoholic fermentation by yeast. Repeatedly, in both methods, indigenous lactic acid bacteria, which are indispensable in the production of *Sake* and *Moto*, grow without being added to the culture. In other words, these lactic acid bacteria are originating from the external environments.

### Impacts of *Kuratsuki* bacteria and adventitious microbes from the external environments

The microorganisms that inhibit surrounding the fermentation reactor, or more precisely, in the breweries, are described as *Kuratsuki* in Japanese. Massive efforts have been made to investigate *Kuratsuki* microbes. One of the significant findings is the study isolating the genus *Kocuria* from sake in multiple batches from the same *Sake* manufacturer in Japan (Terasaki & Nishida 2020). The study also found that *Kocuria* was the unique genus common to all the studied breweries and concluded that the isolated *Kocuria* were *Kuratsuki* microbes. In addition, several *Kuratsuki* microbes have also been isolated, including *Staphylococcus* sp. and *Bacillus* sp., as well as lactic acid bacteria considered necessary for *Sake* production (Terasaki & Nishida 2020; Kanamoto et al. 2021; Takahashi et al. 2021). As an alternative method to identify *Kuratsuki* microbes, a study using amplicon sequencing was reported in 2014 (Bokulich et al. 2014). A comparison between *Kimoto*-style *Moto* and microbes on the surfaces of the brewery was conducted by 16S rRNA and ITS amplicon sequencing. These authors showed that genera present on the surfaces of fermentation equipment surfaces were also detected in *Moto*.

### Tracking the microbiomes from the external environment of *Moto* and their dynamics during the fermentation process

To characterize the source of *Kuratsuki* microbes more deeply, we conducted 16S rRNA amplicon sequencing at *Tsuchida Sake* Brewery in Gunma Prefecture, Japan. We collected the samples from floors, raw materials, and equipment surfaces such as a fermentation tank and a wood stand for growing *Koji*. This brewery produces yeast-free and bacteria-free *Moto*, which means external microbiomes initiate the preparation of *Moto*. We reported their unique microbial transition that *Lactobacillus* did not become dominant in the fermentation starter (Ito et al. 2022). We utilized this sequence data of *Moto* and compared microbial communities between the built environments and the starter culture by counting shared amplicon sequencing variants (ASVs), showing identical sequences in the V3-V4 region of 16S rRNA. Our results suggest that adventitious microbes from the built environment create the initial microbial status, and *Kuratsuki* microbes giving successive fermentation, are screened during competitions in the fermentation.

## 2. Materials and methods

### Sample collection

Samples were collected from 10 locations, including floor, raw materials, and equipment surfaces in the brewery, by a swabbing method (Figure 1). All samples were collected at *Tsuchida Sake* Brewing Company (Gunma, Japan) between October and November 2021. Sterile cotton-tipped swabs (Isohelix Swab: SK-3, Cell Projects Ltd, Harrietsham, UK) were moistened with lysis buffers (BuccalFix Stabilization and Lysis Buffers: BFX, Cell Projects Ltd, Harrietsham, UK) and streaked across an area of the target surface (A square of 20 cm in length and width) for 3 minutes. Samples were immediately frozen and stored until DNA extraction. We collected two samples from different locations on the same sampling targets except for water and used them for library preparation, respectively. All equipment surfaces were swabbed before fermentation. A map and pictures of each floor and equipment are shown in Supplementary Figure 1. The description of the roles of each equipment is listed in Table 1.

**Figure 1.**
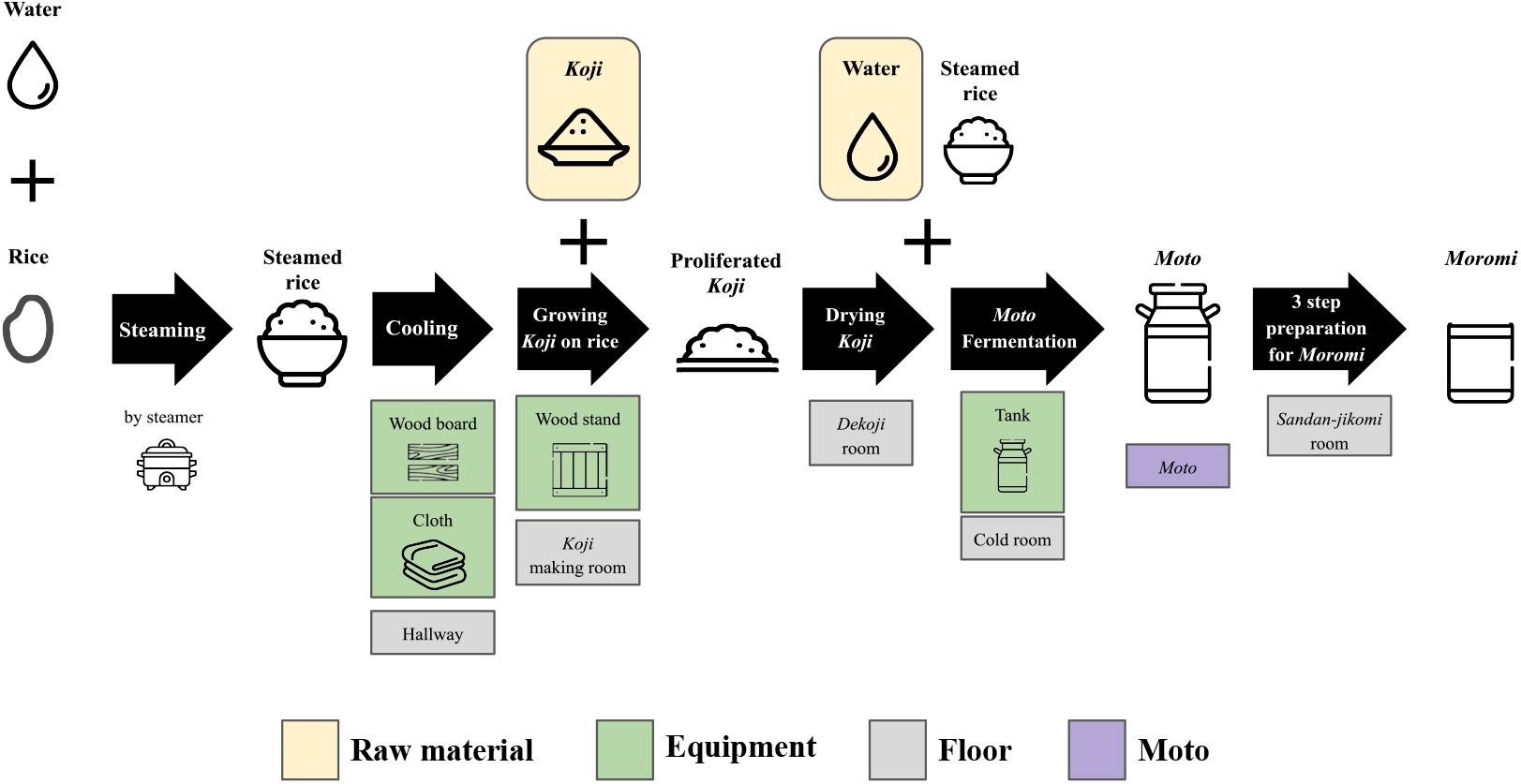
An overview of multiple processes for *Sake* brewing with sampling targets in the study. The group of *Floor* includes the brewery’s floor samples. The group of *Raw materials* includes water and *Koji*. The group of *Equipments* includes the wood board, wood stand, cloth, and tank for *Moto* Fermentation. 10 samples with grid lines were collected in this study. Sample collection was performed in duplicate. A fungus called *Aspergillus oryzae*, which is capable of secreting amylases, is cultivated on steamed rice to create *Koji*. This proliferated *Koji*, along with steamed rice and water, is then placed in an open-top tank to initiate the fermentation process and produce *Moto*. The starter is subsequently mixed with additional *Koji*, steamed rice, and water, and fermentation for a period of 3-5 weeks is done to make *Moromi* through three-step preparation (*Sandan-jikomi*). *Moromi* is separated into *Sake* and spent rice by a filter press.

**Table 1.** The roles of each piece of equipment surveyed in this study.

| **Equipment** | **Roles** |
| --- | --- |
| Wood board | For cooling down the steamed rice |
| Wood stand | For proliferating Koji |
| Cloth | For cooling down the steamed rice |
| Tank | For fermenting *Moto* |

### Total DNA extraction and high-throughput sequencing

Samples were subjected to 750 μl of lysis buffer supplied with the GenFind V2 DNA extraction kit (Beckman Coulter, Indianapolis, U.S.). The suspension was vortexed for 10 min, heat-treated at 100 °C for 10 min, and centrifuged for 5 min at 15,000 rpm. The supernatant was mixed with E.Z. beads (AMR, Tokyo, Japan). DNA fragmentation was performed on an MM-400 (Retsch, Haan, Germany) at a maximum speed of 3 min. The rest of the DNA purification process was performed using a GenFind v2 DNA extraction kit (Beckman Coulter, Indianapolis, U.S.) following the manufacturer’s protocol. DNA was eluted with 80 μl of water. Primers 341F (5’-TCGTCGGCAGCGTCAGATGTGTATAAGAGAGACACCTACGGGNGGCWGCA G-3’) and 806R (5’-GTCTCGTGGGCTCGGGAGATGTGTATAAGAGACAGGACTACHVGGGTATCT AATCC-3’) were utilized to amplify the V3-V4 region of the 16S rRNA gene, and adapter sequences were added by KAPA HiFi HotStart ReadyMix (Roche, Basel, Switzerland) Thermal conditions for PCR were 95 °C for 3 min; 32 cycles of 95 °C for 30 s, 55 °C for 30 s, and 72 °C for 30 s; and a final extension at 72 °C for 5 min. Secondary PCR was performed with forward and reverse primer sequences designated by Illumina. The thermal condition was set to be the same as the first PCR but for 8 cycles. After AMpure XP purification (Beckman Coulter, Indianapolis, U.S.), the libraries were sequenced on the Illumina MiSeq platform (Illumina, Inc., San Diego, USA) by paired-end sequencing of 300 bp. Preparation of DNA samples and libraries and amplicon sequencing were performed at GenomeRead Inc. in Kagawa, Japan.

### Microbiome analysis

Microbiome analysis was performed as previously reported (Ito et al. 2022). Briefly, raw FASTQ files were imported into the QIIME2 platform as qza files. Denoising and quality control of reads was performed by qiime dada2 denoise-paired, and reads were classified into amplicon sequencing variants (ASVs) (Callahan et al. 2016). The SILVA database SSU 138 was utilized with qiime feature-classifier classify-sklearn for taxonomic assignment (Bokulich et al. 2018; Quast et al. 2013). ASVs classified as chloroplast, mitochondria, and unassigned were excluded from statistical analysis. Although the genus *Latilactobacillus* is a taxon that was recently derived from the genus *Lactobacillus* (Zheng et al. 2020), the SILVA database SSU 138 did not include this reclassification of the genus. Thus, the conventional taxonomic name *Lactobacillus* was adopted in this study. One sample from the cold room floor did not have enough sequencing depth and was excluded from the downstream analysis. The depth of sequence reads differed among samples, and we subsampled from each sample to 4,724 reads each to normalize the data. Subsampling is an approach for inferring microbiome differences between samples and has been reported to be a suitable analytical method when analyzing new datasets (Hughes & Hellmann 2005). To evaluate the effect of sequence read counts on microbiome diversity assessment, we examined changes in the value of the number of ASVs over a range of reading counts from 0-8,000 by rarefaction curves. The rarefaction curves of the number of ASVs were leveled off when the number of reads reached approximately 4,000 (Supplementary Figure 2).

### Calculation of shared ASVs

Sequence data obtained in this study were confronted to those previously obtained (PRJDB13924) to detect ASVs present in both datasets (Ito et al. 2022). Shared ASVs in this study were defined as ASVs present at more than 1% were considered to detect those shared by different types of data sets. The calculation was conducted by R version 4.2.1 and Phyloseq (McMurdie & Holmes 2013) version 1.40.0.

### Statistical analysis

Kruskal–Wallis tests were used to compare the relative abundances among sample types. Mann–Whitney U tests were used to compare all combinations of elements within the sample types. All multiple testing corrections were performed by computing FDRs using the Benjamini-Hochberg method, and adjusted P values < 0.05 were considered statistically significant. Differentially abundant genera between groups were identified using linear discriminant analysis (LDA) effect size (LEfSe) analysis (Segata et al. 2011). Only bacterial genera reaching the LDA score threshold of 3.0 and with a p value lower than 0.01 are shown. Data visualization was performed by R version 4.2.1, the ggplot2 package and ggprism (Dawson 2022).

## 3. Results

### The phylum-level bacterial profiles in the *Sake* brewery

The number of reads after DADA2-based denoising is described in Supplementary Table 1. The relative abundances of the top 7 phyla are shown as a heatmap (Figure 2A). The bacterial profiles of floor surfaces showed relatively high abundances of *Bacteroidota, Proteobacteria, Firmicutes*, and *Actinobacteria*. Equipment surfaces (wood board, wood stand, cloth, tank) showed similar trends as floor surfaces, but cloth and tank showed lower relative abundance in *Bacteroidota* and higher relative abundance in *Deinococcota*. Raw materials (*Koji* and water) are rich with *Firmicutes* and *Proteobacteria*. The relative abundances of the four phyla were significantly different among the three sample types (Figure 2B-E). *Firmicutes* had the highest median of over 30% for raw materials, and Mann–Whitney U tests also showed a significant difference in raw materials against floor surfaces (p-value = 0.05) and equipment surfaces (p-value = 0.016) (Figure 2B). Otherwise, floor surfaces and equipment surfaces showed higher relative abundances in *Bacteroidota*, *Deinococcota*, and *Actinobacteria* (Figure 2C-E).

**Figure 2.**
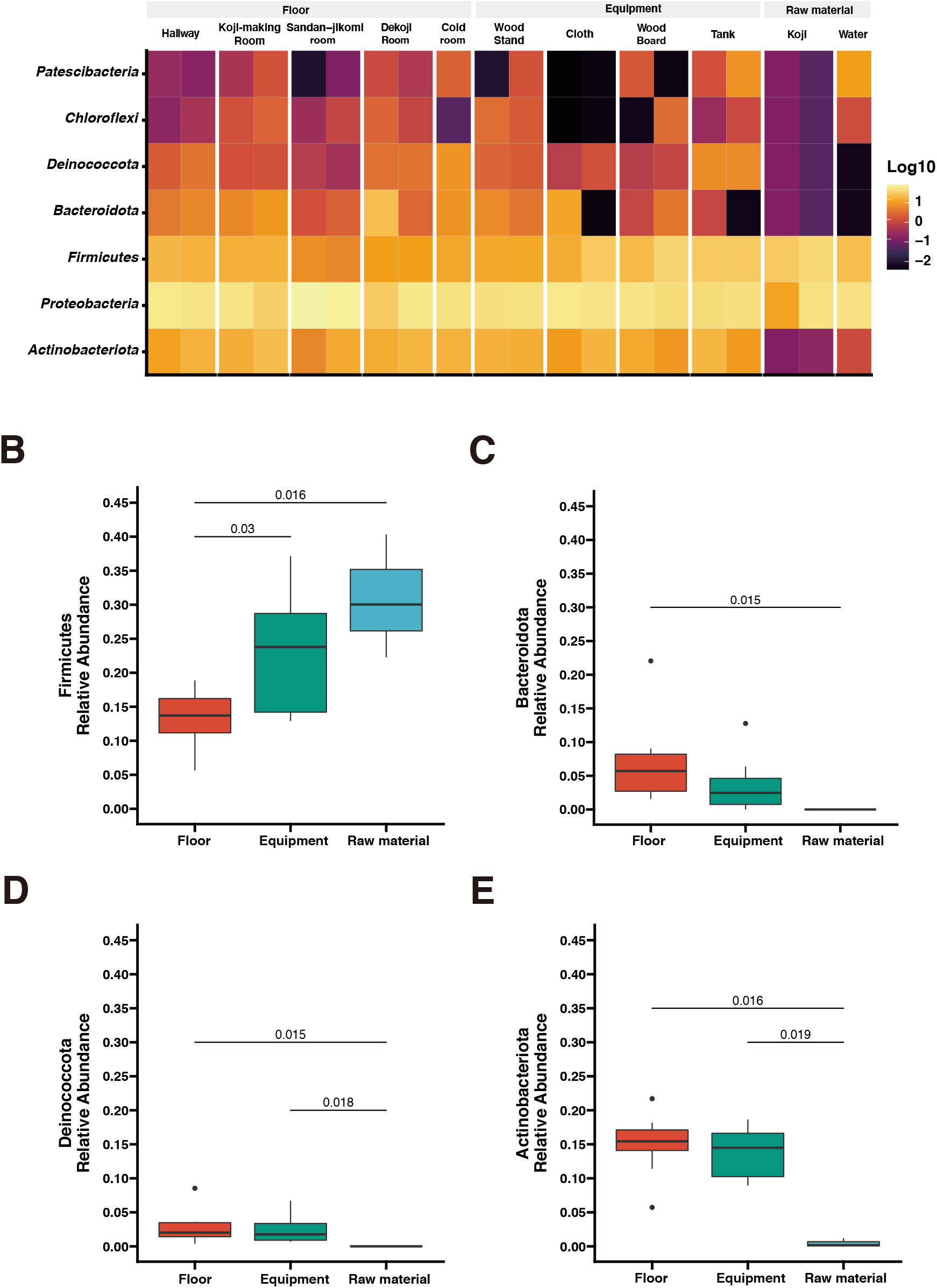
(A) Taxon abundance heatmap at the phylum level. Heatmap depicts the relative abundances (log10 scale) of the top 7 phyla across all surfaces. (B-E) Box plot representation for phylum-level comparison among each group. Four phyla (*Firmicutes, Bacteroidota, Deinococcota, Actinobacteriota*) were compared across groups in the brewery. Statistical analysis was computed using Mann-*U* tests.

### Distribution patterns of bacterial genera associated with *Kimoto*-style fermentation

Although the genus *Latilactobacillus* and 22 new genera were derived from the genus *Lactobacillus* (Zheng et al. 2020), the classifier used in this study did not include this reclassification of the genus. The top 15 genera are shown in a stacked bar graph (Figure 3); the rest are noted as the *remainder. Sphingomonas* was widely detected with a high abundance ratio within samples grouped in floor surfaces (4.2~33.5%). 28.3% and 27.5% *Halomonas* were detected from the *Sandan-jikomi* (three-step preparation for *Moromi*, fermentation mash) room floor. *Koji* had high abundances in *Anaerobacillus* (30% and 8.7%, respectively) and *Staphylococcus* (20.1% and 4.7%, respectively). The heatmaps sorted by *Firmicutes*, *Bacteroidota*, *Deinococcota*, and *Actinobacteria* are shown in Figure 3B-E to identify the unique genera within each sample. The relative abundance of the genus *Lactobacillus*, considered to be necessary for brewing, was found in more than 1% of only one sample of the cold room floor (3.6%) and the wood board (2.8%) (Supplementary Figure 3B). Both surface types *Leuconostoc*, the dominant species of *Moto* in the brewery, was found in the tank, *Koji* making room floor, *Dekoji* room floor, and cold room floor (Supplementary Figure 3B). *Bacillus* was widely detected in water (8.7% and 1.1%, respectively), wood board (8.3% and 0%, respectively), and cloth (8.8%, and 1.7%, respectively) (Supplementary Figure 3B). LFfSe analysis identified *Anaerobacillus, Methylobacterium, Escherichia*, and *Delftia* as differentially abundant in raw materials compared with other sample types (Supplementary Figure 4).

**Figure 3.**
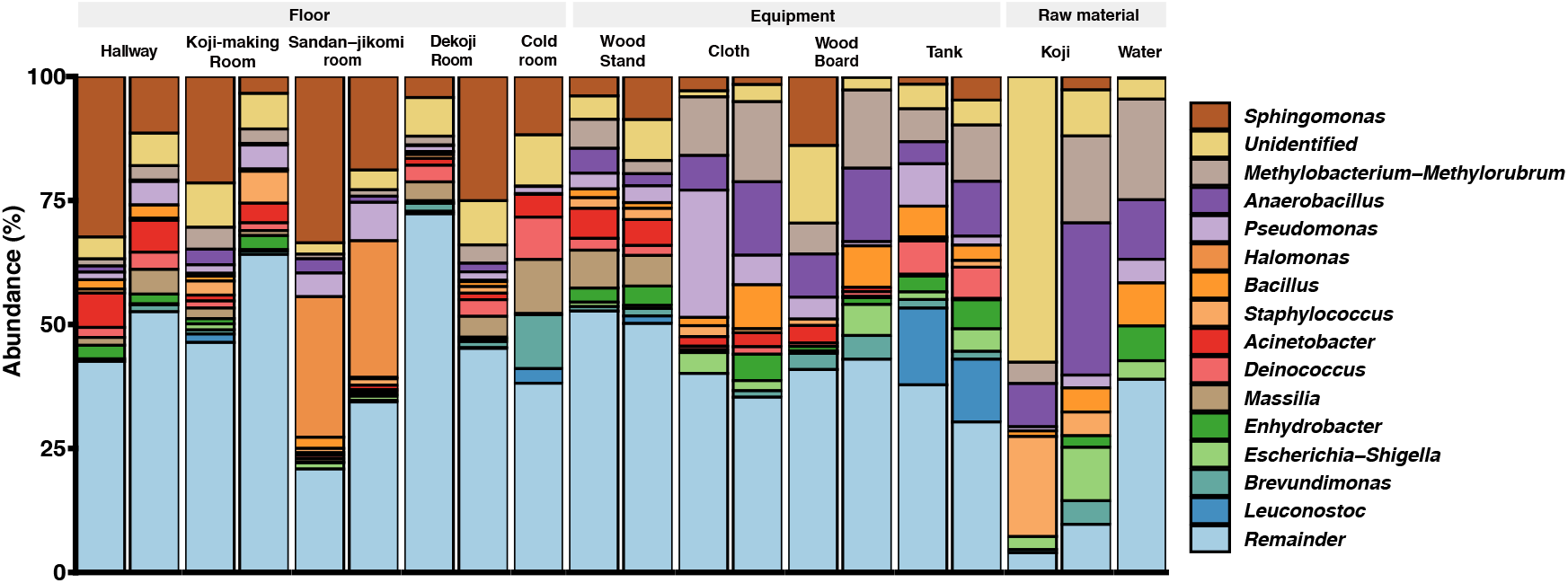
Genus-level taxonomic composition across the environmental samples.

### Changes in the relative abundances of microbes from the external environment during *Moto* fermentation

Principal coordinates analysis (PCoA) based on unweighted UniFrac distances was performed (Figure 4A). The dots were divided into three major groups according to their sample types. Raw materials and equipment surfaces are located between *Moto* and the group of floor surfaces. Floor surface samples differ substantially from *Moto*.

**Figure 4.**
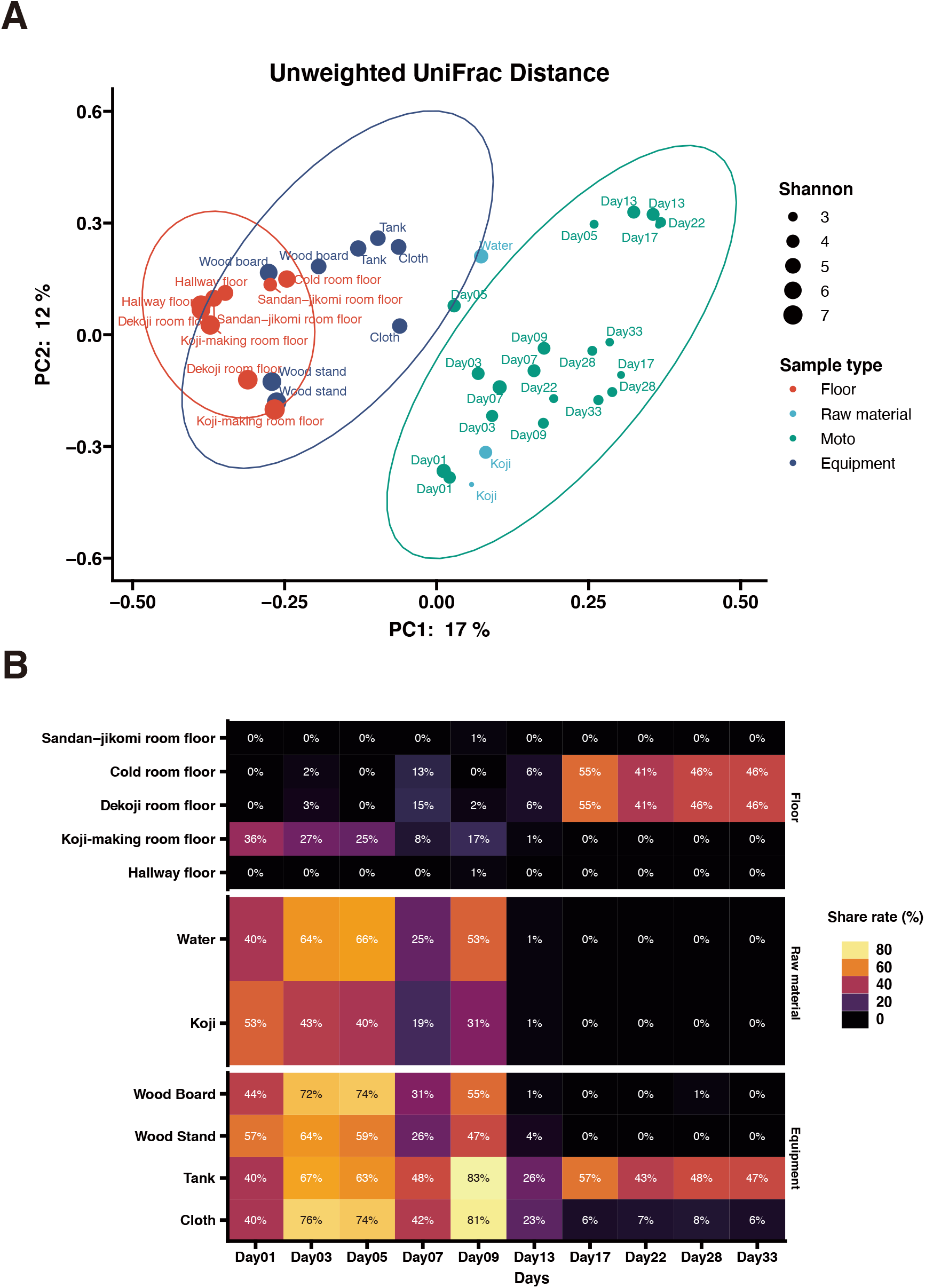
(A) Principal-coordinate analysis (PCoA) by Weighted UniFrac distance of environmental samples and *Moto* samples. The dot size shows the values of Shannon diversity of each sample. (B) Time-series changes in the relative abundance of shared ASVs in *Moto* during the *Kimoto*-style fermentation period. More than 40% of the microbiomes in *Moto* on Day 33 shared the ASVs with the tank surface, cold room floor, and *Koji*-making room floor samples.

The results of the shared ASV analysis are shown as a heatmap (Figure 4B). Bacteria confirmed on the surface of all raw materials and equipment surfaces showed shared ASVs with *Moto* samples throughout Day 1 to Day 9. The *Koji* making room also showed them from Day 1 to Day 5. The ratio of shared ASVs increased on Day 3 and Day 5. The tank and cloth reached 83% and 81%, respectively, on Day 9. Day 13 showed a decrease in the ratio of the tank (26%) and cloth (23%) and an increase in the cold room (6%) and *Dekoji* room (6%). From Day 17 to Day 33, only the tank, *Dekoji* room floor, and cold room floor showed shared ASVs. The tank, cold room, and *Dekoji* room showed more than 40% in the later days. Further details of ASV-assigned genera shared by *Moto* on Day 33 are shown in Table 2. ASVs assigned to *Pseudomonas* in *Moto* were shared between the tank and cloth, and *Serratia* and *Hafnia-Obesumbacterium* were shared by cloth. Tank, cold room, and *Dekoji* room showed *Leuconostoc* sharing with *Moto*. The shared ASV of *Leuconostoc* consisted of a single ASV unit.

**Table 2.** The taxon of shared ASV (Genus-level) between *Moto* on Day33 and other environments.

| Taxon | Confidence | Shared surface types |
| --- | --- | --- |
| *Pseudomonas* | 0.999 | Tank, Cloth |
| *Leuconostoc* | 0.999 | Tank, Koji room floor, Cold room floor |
| *Serratia* | 0.904 | Cloth |
| *Hafnia-Obesumbacterium* | 0.858 | Cloth |

## 4. Discussion

### Bacterial profiles differed by sample types

We found different phyla profiles between the raw materials and environmental samples (floor surfaces and equipment surfaces). The microbial composition of floor surface samples were diverse compared with the rest of the two sample types (Figure 3, Figure 4A). Previous studies indicate that floors in indoor environments are known to be rich in microbes (Hospodsky et al. 2012; Ruiz-Calderon et al. 2016). For this reason, we included floor samples rather than walls and ceilings. The built environments are also complex structures, and microbial communities differ by conditions such as surface materials and the possibility of human interaction. In fermentation starter production, the manufacturer inputs multiple ingredients and uses equipment surfaces. Also, *Moto* is produced in multiple rooms based on the steps of production. These direct interactions with sources of microbes can make microbiomes in fermentation starters of early days exceedingly divergent. *Firmicutes* are detected in high abundance on dry (e.g., floors, cabinets, microwaves) and cold (e.g., inside of the refrigerators) surfaces in the kitchen (Flores et al. 2013) and refrigerator surfaces (Einson et al. 2018), but in this study, they were most abundant in raw materials (Figure 2B).

Genus-level composition (Figure 3A) shows that *Koji* samples have *Bacillus, Anaerobacillus, Staphylococcus*, and *Sphingomonas*. These genera were confirmed in *Sokujo*-style *Moto* by Sanger sequencing of their 16S rRNA sequences in the previous report (Terasaki et al. 2017). In this study, these genera were especially detected in the architectural samples. As other features, *Koji* is made by bare hands; therefore, it is considered that *Staphylococcus*, a genus encompassing species known to be present on skin (Cogen et al. 2008), may have been contaminated in the *Koji* and the floors.

We reported that *Leuconostoc* was dominant in the fermentation of *Moto*, and the switch of the predominant lactic acid bacteria from *Leuconostoc* to *Lactobacillus* was not observed in the fermentation process of this brewery (Ito et al. 2022). Several studies reported that *Lactobacillus* was detected in tanks (Bokulich et al. 2015; Bokulich et al. 2014) that can be directly in contact with raw materials and *Moto*. However, *Lactobacillus* was detected only on the cold room floor and the wood board. In contrast, *Moto* did not contact these sites. However, *Leuconostoc* was detected in the tank of *Moto*. This result suggests that appreciable concentrations of *Leuconostoc* are introduced from the tank to the fermentation reaction of *Moto*. This consideration is consistent with findings in a previous report (Bokulich et al. 2014) that surface contact is the most significant predictor of bacterial composition in the *Sake* brewery. We also concluded that the sources of *Kuratsuki* microbes are possibly surfaces of equipment surfaces and raw materials that have high chances of direct contact with *Moto*. Therefore, the *Moto* in *Tsuchida Sake* Brewery may not have shown the lactic acid bacteria switch from *Leuconostoc* to *Lactobacillus* in the fermentation process due to the lack of the sources of *Lactobacillus*.

### Shared ASV analysis revealed the interaction between environmental samples and *Moto*

In the PCoA of unweighted UniFrac distances (Figure 4A), raw materials are located near the fermentation starter because they are an input of Moto. Next are the equipment surfaces that are in direct contact with *Moto*. The floor is not in direct contact with the fermentation starter and thus differs substantially from the group structure of *Moto*. In this study, the floor was employed as the floor surfaces group. This is based on our expectation that microbes may fall on the floor during manufacture. However, experimental data indicate that surfaces that can be in direct contact may have a more significant impact on *Moto*. The bacterial profile of brewing equipment surfaces has yet to be fully revealed, and further research is needed.

In this study, we investigated how bacteria are transferred to *Moto* as a source. Bacteria on the equipment surfaces and floor surfaces were confirmed in the early stage of *Moto* (Figure 4B). Thus, the shared ASVs results indicate that *Leuconostoc*, which plays an essential role in *Moto*, was screened within adventitious microbes and eventually became the dominant species.

### Technical limitations in 16 rRNA amplicon sequencing

Because of the limited classification resolution of 16S rRNA amplicon sequencing, it is challenging to rigorously predict microbial sharing between fermentation starters and the environment. Calculating shared ASVs or statistical methods is a convenient way to track microbial sources. However, high-resolution sequencing, such as metagenome sequencing, must be performed to clarify precisely which places or sources in bacteria affect fermentation. Lineages of bacteria between two groups can be compared by metagenome-assembled genomes (MAGs) and show more precision in tracking at the level of whole chromosomal sequences, not just the restricted regions of 16S rRNA.

We reported that the ethanol concentration increased as fermentation progressed (Ito et al. 2022). According to ethanol production from yeast, bacteria should decrease gradually. However, 16S rRNA amplicon sequencing is a DNA-based detection method, and it is impossible to discern between viable and nonviable bacteria. In fermented foods, viable bacteria can be considered to have a significant influence, and conventional microbiology experiments or flow cytometry with staining is required to verify the viable cells. In addition, fungi also play a pivotal role in *Sake* fermentation. Therefore, metagenome sequencing is necessary to unveil the interaction between bacteria and fungi, especially for fermentation utilizing *Koji, Aspergillus oryzae*, and *Saccharomyces cerevisiae*.

Also, microbes in the built environments and equipment are not as abundant as in fermented food. *Kuratsuki* microbes inhabit breweries in a heterogeneous manner. In our samples, we confirmed the variety of relative abundances in some genera between duplicated experiments (Figure 3). We sampled each duplicate set from different locations. This can explain by biases in library preparation, including PCR or sampling bias in each experiment. Multiple sampling points in the exact locations can give higher resolutions in rare ASVs, and shared ASVs analysis can show precision results.

### *Kuratsuki* microbiota as a community structure and their impact on sake brewing

Tracing the dynamic network of microbial communities further contributes to understanding this traditional liquor. According to the previous study, microbial diversity decreases with the number of fermentation days of *Moto* (Bokulich et al. 2014). Another study performing microbiome analysis of *Moto* from five different *Sake* breweries reported that the bacterial communities of *Moto* were very diverse, but that diversity decreased over time through multiple microbial transition patterns (Takahashi et al. 2021). It was also reported that ethanol produced by yeast inhibits the growth of adventitious microbes (Terasaki & Nishida 2020). Microbes produce metabolites that influence microbial dynamics over time. Both metabolome and microbiota interact mutually. Profiling the metabolome of *Sake* breweries can deepen the understanding of the roles of *Kuratsuki* microbes.

Some strains have been isolated from *Moto* and *Sake*. *Latilactobacillus sakei* was isolated from *yamahai-moto* (Tsuji et al. 2018). *Lactiplantibacillus plantarum* was also isolated from different *yamahai-moto*. These lactic acid bacteria were not confirmed in our study but they can also be *Kuratsuki* microbes and further study in different breweries should be conducted.

Furthermore, some adventitious microbes have been found to possess activity interfering with the growth of bacteria that spoil *Sake*, potentially benefiting *Sake* brewing (Taniguchi et al. 2010) or producing beneficial chemical components that contribute to the flavor, taste, and quality of *Sake* (Akaike et al. 2020).

Mapping the microbiomes in *Sake* breweries may also be important in terms of quality and safety in *Sake* production (De Filippis et al. 2021). In the past, studies have yet comprehensively examined changes in microbes sharing between different sample types (floor surfaces and equipment surfaces) and *Moto* in a brewery during the fermentation process with a lens of amplicon sequencing. Previous studies have isolated *Kocuria* and *Bacillus* as viable microbes (Kanamoto et al. 2021; Terasaki et al. 2021), indicating that certain microbes can live in *Moto*, such as lactic acid bacteria and yeast, and may be involved in *Sake* brewing. The possibility of horizontal gene transfer among microorganisms in *Sake* breweries (Terasaki et al. 2021) has also been demonstrated; instead of targeting only specific bacteria, such as lactic acid bacteria, the community-level structure should be targeted as *Kuratsuki* microbiota. Isolation of viable microbes and whole genome sequencing can give a deep insight into gene transfer.

Finally, understanding microbial diversity and dynamics are about preserving traditional food culture and food production safety. Sake requires attention to spoilage, such as microbial contamination and spoilage, which can cause spoilage and may be helpful for safety and quality control. This research can be an essential milestone in unraveling the mysteries of fermentation phenomena. Because each brewery produces different flavors of sake with different ingredients and equipment surfaces, it is necessary to further research to profile microbiomes in sake breweries. Additionally, these contributions can lead to process control, spoilage prevention, food safety, and the protection of traditional cultures.

## Data Availability Statement

The datasets generated for this study can be found in National Center for Biotechnology Information (NCBI) Sequence Read Archive (SRA) under an accession number DRR423705 (https://www.ncbi.nlm.nih.gov/sra/?term=DRR423705).

## Author Contributions

K.I. and R.N. conceived this study. K.I. and K.K. designed the experiments. K.I. and R.N. wrote the original draft. R.N. performed microbiome analysis. K.I. and R.N. performed the statistical analysis. Y.T., G.H., and K.K. collected samples and provided the records required for discussion. T.N. supervised the study and edited the manuscript. All authors contributed to the article and approved the submitted version.

## Acknowledgments

Samples were collected with the help of members of the *Tsuchida Sake* Brewery. GenomeRead Inc. performed amplicon sequencing. All authors thank Morgenrot Inc. for providing the computational environment for the analysis. R.N. is a graduate student of the Medical Innovation Program at Kyoto University and supported by the JST SPRING program, Grant Number JPMJSP2110. We cited icons for Figure 1 from https://www.flaticon.com/free-icon/.

