## Supplemental files for "A dark matter in *sake* brewing: Origin of microbes producing a *Kimoto*-style fermentation starter"

[†] **Equal contribution**

**This file includes:**

Supplementary figure 1-4

Supplementary Table 1

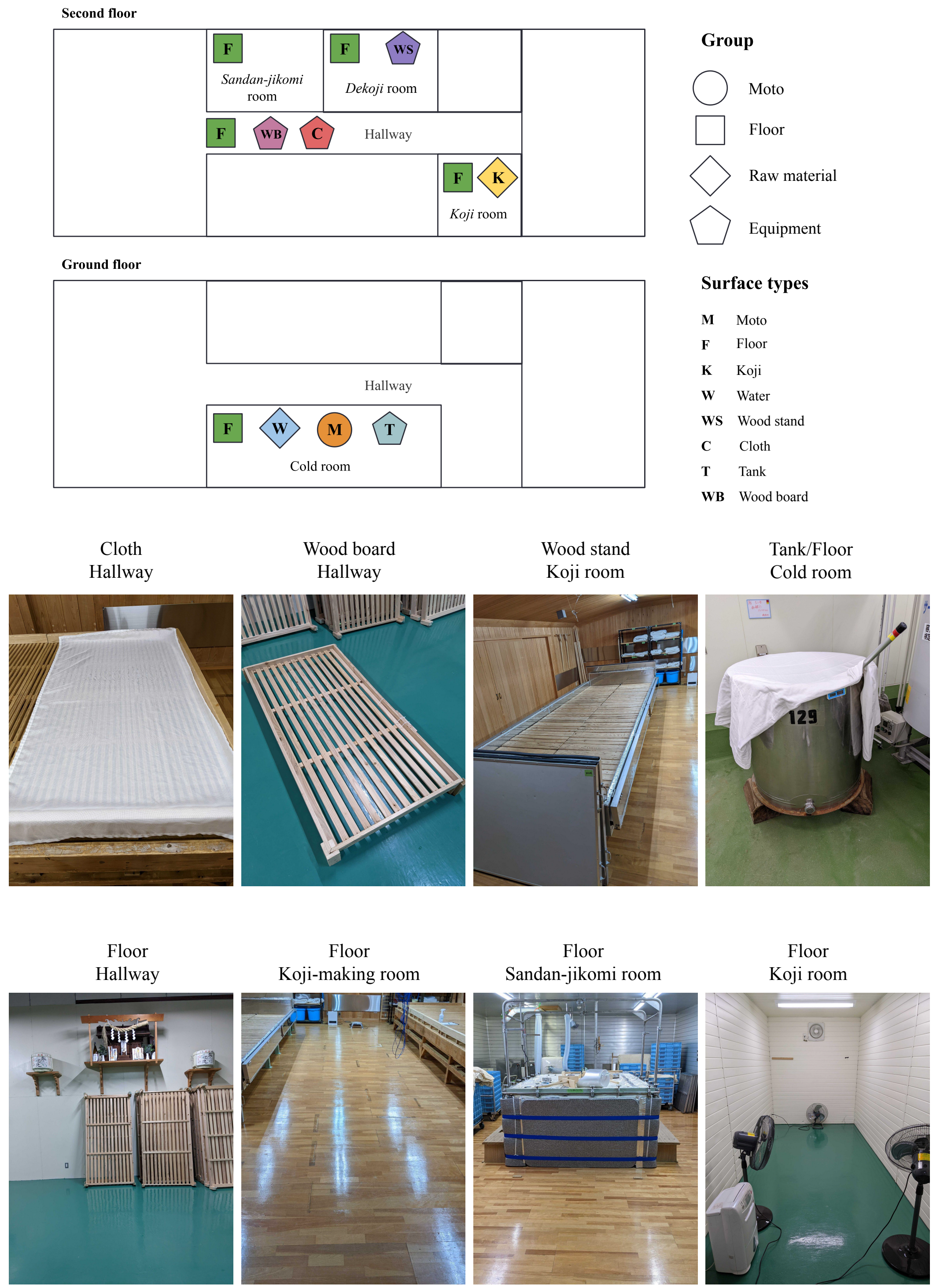


**Supplementary Figure 1.** A map of the brewery surveyed in this study. Shapes of the dots describe groups of samples. Pictures of equipment and floor are shown below the map.

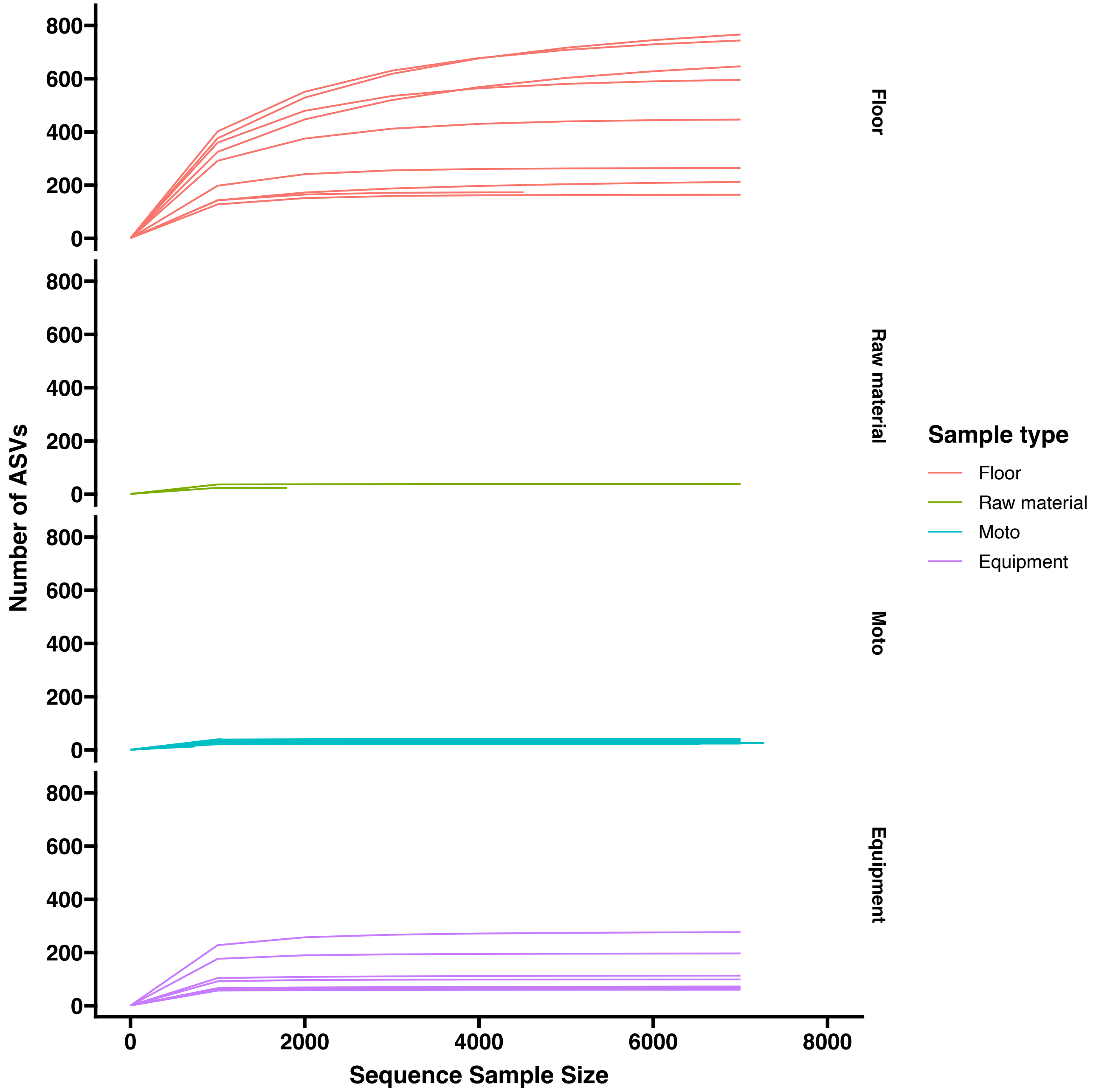


**Supplementary Figure 2.** A rarefaction curve showing the accumulation of the number of ASVs by sampling depth.

A

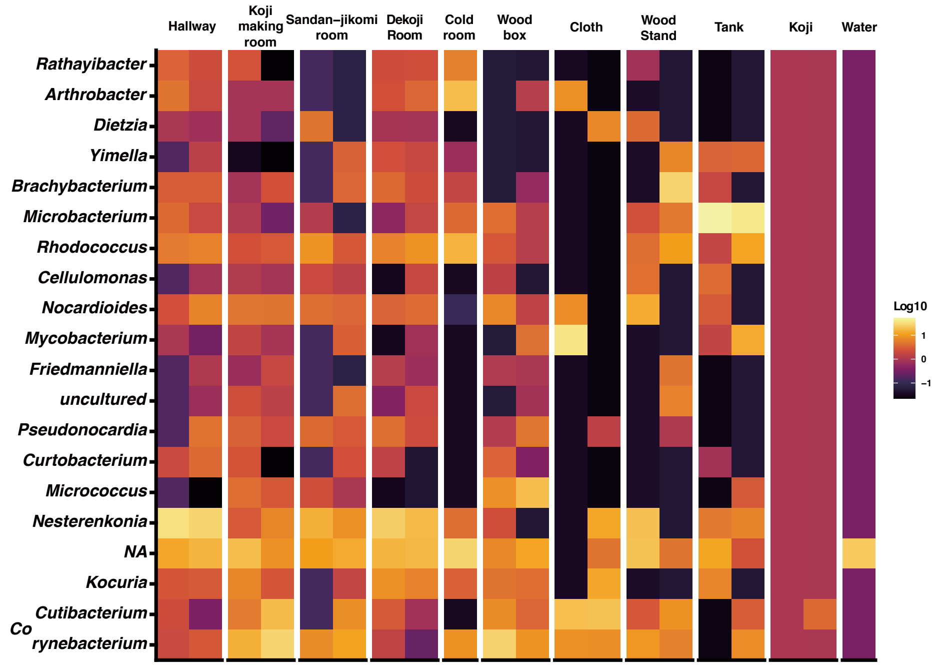
Hallway
Koji making room
Sandan–jikomi room
Dekoji Room
Cold room
Wood box
Cloth
Wood Stand
Tank
Koji
Water
Rathayibacter
Arthrobacter
Dietzia
Yimella
Brachybacterium
Microbacterium
Rhodococcus
Cellulomonas
Nocardioides
Mycobacterium
Friedmanniella
uncultured
Pseudonocardia
Curtobacterium
Micrococcus
Nesterenkonia
NA
Kocuria
Cutibacterium
Co rynebacterium
Log10
1
0
−1

B

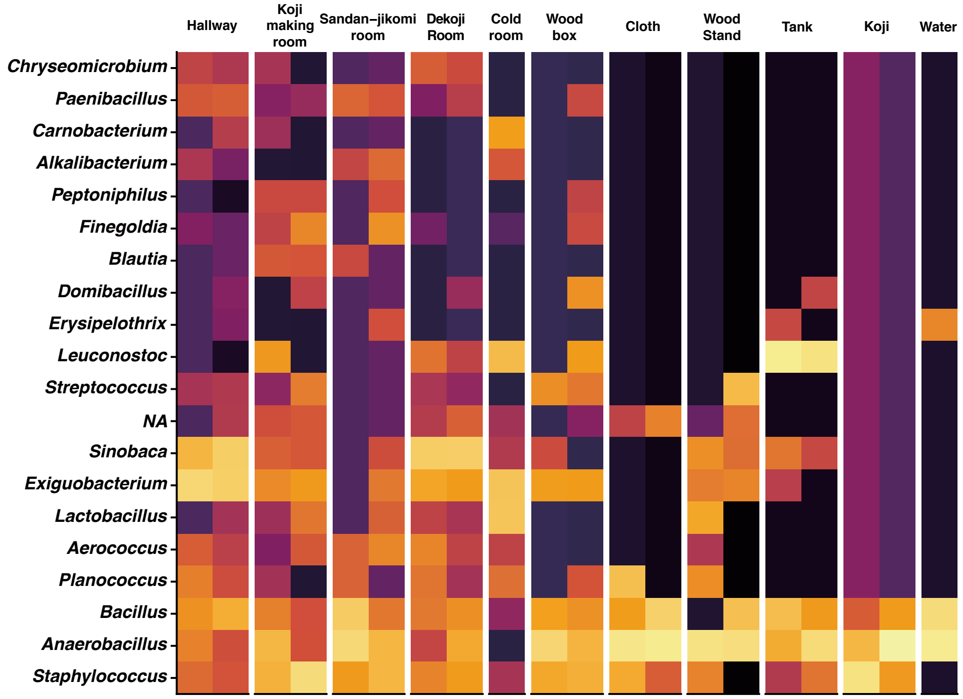
Hallway
Koji making room
Sandan–jikomi room
Dekoji Room
Cold room
Wood box
Cloth
Wood Stand
Tank
Koji
Water
Chryseomicrobium
Paenibacillus
Carnobacterium
Alkalibacterium
Peptoniphilus
Finegoldia
Blautia
Domibacillus
Erysipelothrix
Leuconostoc
Streptococcus
NA
Sinobaca
Exiguobacterium
Lactobacillus
Aerococcus
Planococcus
Bacillus
Anaerobacillus
Staphylococcus

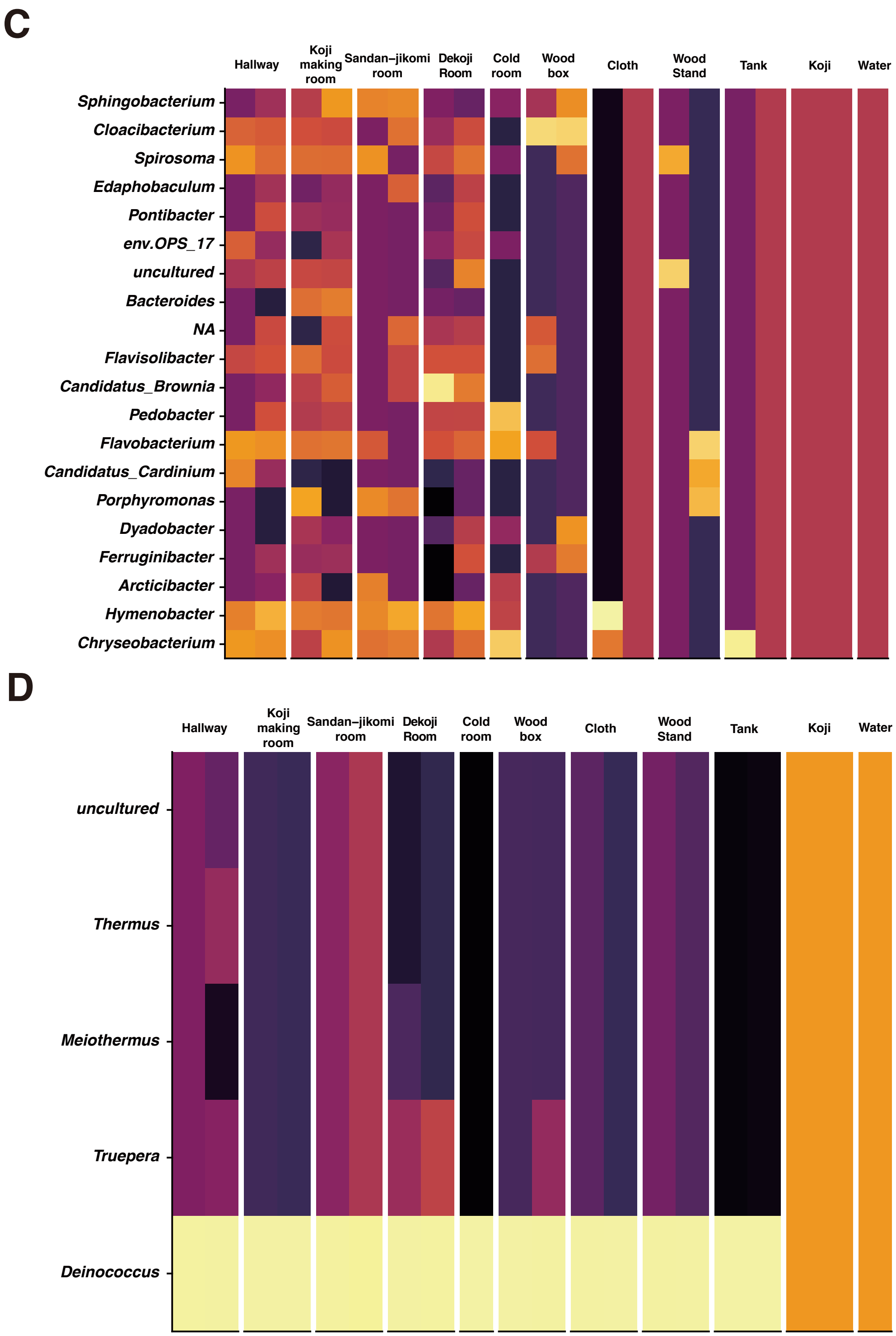


**Supplementary Figure 3.** Heatmaps showing the relative abundances (log10 scale) at the genus level of 4 phyla (*Firmicutes, Bacteroidota, Deinococcota, Actinobacteroidota*).

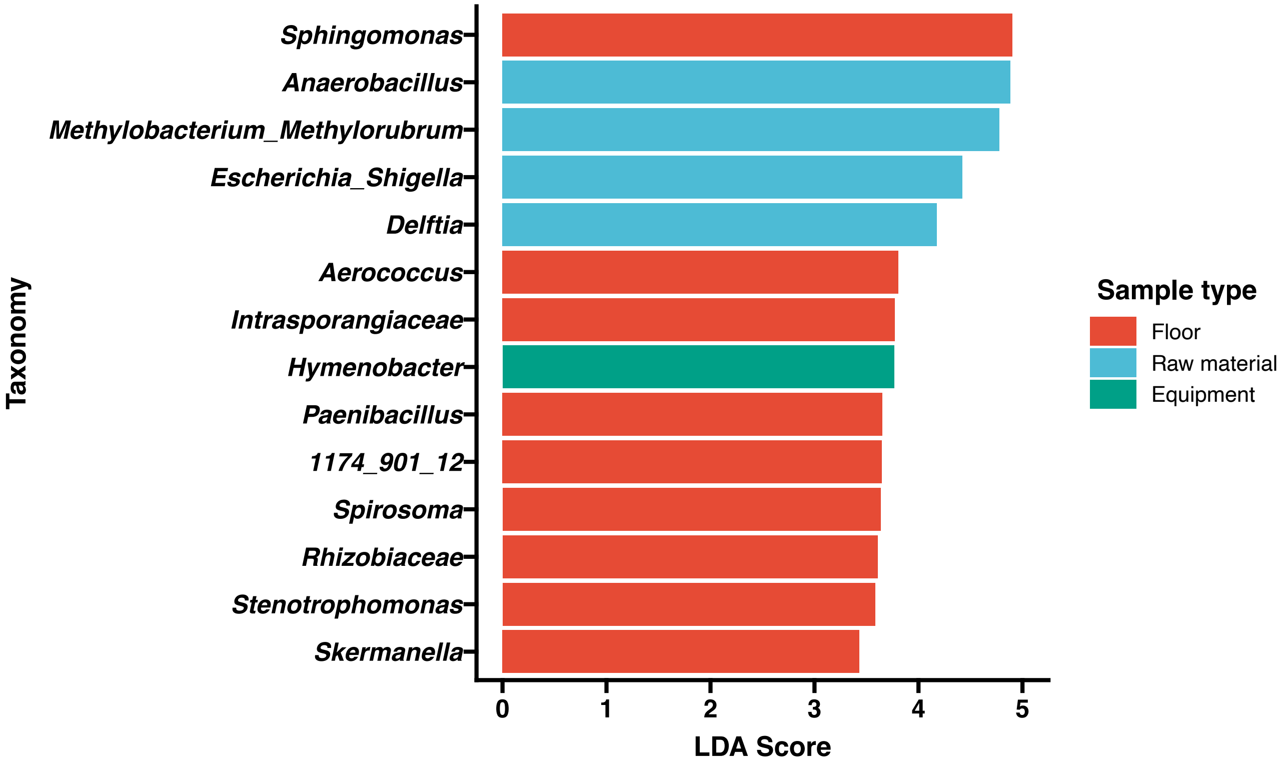


**Supplementary Figure 4.** Differential abundance microbes in the microbial communities by sample types from LEfSe computation.

**Supplementary Table 1.** A table of the number of reads per sample after the DADA2 denoising step.

| sample-id | Sampling place | Reads |
|---|---|---|
| S01 | cloth | 27211 |
| S02 | cloth | 21495 |
| S03 | wood stand | 18052 |
| S04 | stand stand | 21057 |
| S05 | wood board | 13138 |
| S06 | wood board | 15223 |
| S07 | tank | 19977 |
| S08 | tank | 19367 |
| S09 | water | 17010 |
| S10 | koji | 14918 |
| S11 | koji | 17240 |
| S12 | Hallway | 4724 |
| S13 | Hallway | 27393 |
| S14 | Koji room | 20794 |
| S15 | Koji room | 29908 |
| S16 | Sandan-jikomi room | 11027 |
| S17 | Sandan-jikomi room | 9334 |
| S18 | Dekoji room | 22469 |
| S19 | Dekoji room | 11017 |
| S20 | Cold room | 0 |
| S21 | Cold room | 15978 |
